# Systems Pharmacology Reveals Type I Interferon and Myeloid-Like B Cell Reprogramming as Druggable Axes in Antiphospholipid Syndrome

**DOI:** 10.64898/2026.04.28.721277

**Authors:** Bin Sun, Yiqiao Lu, Wei Liu, Chengze Wang

**Affiliations:** Jiangnan University Medical Center, Wuxi, China; Women’s Hospital School of Medicine Zhejiang University, Zhejiang University, Hangzhou, China; Zhejiang University School of Medicine, Clinical Research Center for Oral Diseases of Zhejiang Province, Key Laboratory of Oral Biomedical Research of Zhejiang Province, Cancer Center of Zhejiang University, Hangzhou, China

**Keywords:** antiphospholipid syndrome, systems pharmacology, drug repurposing, WGCNA, single-cell RNA sequencing, Connectivity Map, network pharmacology, molecular docking, precision medicine, interferon signaling, B cell reprogramming

## Abstract

Antiphospholipid syndrome (APS) lacks targeted therapies beyond anticoagulation, and its molecular heterogeneity remains poorly characterized. We employed an integrative systems pharmacology approach combining weighted gene co-expression network analysis (WGCNA), single-cell RNA sequencing, Connectivity Map (CMap) screening, and molecular docking to identify druggable targets in APS. WGCNA of bulk RNA-seq data from neutrophils (n = 18) and whole blood (n = 88) identified two disease-associated modules: ME10 (176 genes, r = 0.77, interferon-I signaling) and ME2 (3409 genes, r = 0.79, degranulation/innate activation). Single-cell analysis of 26,936 B cells revealed transitional B cells with elevated ME2 scores and aberrant SPI1 expression, suggesting myeloid-like transcriptional reprogramming. CMap analysis ranked chloroquine, a first-line APS therapy, among top ME2 candidates (NCS = −2.07), validating the computational approach. DrugBank mapping identified 14 FDA-approved drugs targeting module genes, and a 3-gene machine learning signature (CORO1A, ANKRD22, IFITM1) achieved cross-tissue validation AUC of 0.802. External validation confirmed ME2 pathway modulation by NAPc2 intervention and cross-tissue module conservation in platelets. Patient-level ME10 x ME2 stratification revealed four molecular subtypes with distinct pathway activation profiles. This framework nominates druggable targets across both IFN-I and degranulation pathways, providing a foundation for pathway-guided precision medicine in APS.

## Introduction

Antiphospholipid syndrome (APS) is a systemic autoimmune disorder characterized by arterial and venous thrombosis, recurrent pregnancy loss, and the persistent presence of antiphospholipid antibodies (aPL).[1–3] Despite affecting approximately 1-5% of the general population and representing a major cause of acquired thrombophilia, therapeutic options for APS remain limited to anticoagulation strategies that do not address the underlying pathogenic mechanisms.[4–5] The heterogeneity in clinical manifestations and treatment responses among APS patients suggests distinct molecular endotypes that have yet to be systematically characterized.[6]

Recent advances in systems biology and computational pharmacology have enabled the identification of disease-associated molecular networks and the rational repositioning of FDA-approved drugs for new therapeutic indications.[7–8] Weighted gene co-expression network analysis (WGCNA) has emerged as a powerful approach for identifying co-expressed gene modules that reflect shared biological functions and regulatory mechanisms.[9–10] When integrated with single-cell transcriptomics, WGCNA can reveal cellular heterogeneity and cell-type-specific pathway dysregulation that may be obscured in bulk tissue analyses.[11] Furthermore, the integration of transcriptomic signatures with drug-target databases such as DrugBank and expression-based screening platforms like the Connectivity Map (CMap) enables systematic identification of therapeutic candidates that can reverse disease-associated gene expression patterns.[12–14]

Previous studies have implicated interferon signaling, neutrophil activation, and complement pathways in APS pathogenesis.[15–20] However, these investigations have largely relied on candidate gene approaches or focused on individual cell types, lacking a comprehensive systems-level analysis that integrates multi-omics data to identify druggable targets. Moreover, the molecular basis for patient-to-patient variability in APS remains poorly understood, limiting the development of precision medicine approaches.[21–25]

Recent bioinformatics studies have begun to apply co-expression network approaches specifically to APS transcriptomics. A WGCNA-based analysis of whole blood RNA-seq data from thrombotic APS patients identified five co-expression modules, with STAT1 emerging as a central hub gene and promising pharmacological target through drug-gene interaction analysis.[26] Separately, an integrated bioinformatics and machine learning study employing WGCNA on neutrophil RNA-seq data (GSE102215) uncovered shared transcriptomic biomarkers (CCR1, MNDA, S100A8, CXCL2) between APS and recurrent miscarriage, nominating candidate therapeutic targets.[27] While these studies validate co-expression network analysis as a productive computational strategy in APS, they did not integrate single-cell RNA sequencing to characterize cellular heterogeneity, expression-based drug screening (CMap/LINCS) to identify candidate therapeutics, molecular docking for structural plausibility assessment, or patient-level molecular stratification. The present study addresses these gaps through a comprehensive multi-layer systems pharmacology framework, delivering actionable drug repositioning candidates validated across independent datasets.

In this study, we employed an integrative systems pharmacology framework to: (i) identify core disease-associated gene modules through WGCNA of bulk transcriptomic data from neutrophils and whole blood; (ii) characterize cellular heterogeneity and module-specific functional states using single-cell RNA sequencing of B cells; (iii) stratify patients based on module expression profiles; (iv) perform network-based drug repurposing by integrating DrugBank target predictions and CMap expression-based screening; and (v) validate predicted drug-target interactions through molecular docking and external experimental datasets. Our approach identifies druggable targets across both ME10 (IFN-I) and ME2 (degranulation) pathways, revealing diverse therapeutic opportunities that extend beyond traditional kinase inhibition strategies.

## Materials and Methods

### Data Acquisition and Processing

Three transcriptomic datasets were obtained from GEO: neutrophil bulk RNA-seq (GSE102215; 9 APS, 9 controls), whole blood bulk RNA-seq (GSE205465; 53 APS, 35 controls), and single-cell B cell RNA-seq (GSE262240; 26,936 cells).[28–29] Bulk data were normalized using DESeq2 (v1.38.0),[28] with variance-stabilizing transformation for downstream analyses. Single-cell data were processed with Seurat (v5.0.0),[29] normalized by SCTransform,[30] batch-corrected using Harmony,[31] and clustered via the Louvain algorithm. External validation datasets included GSE252972 (in vitro NAPc2 intervention), GSE252397 (mouse splenic DC), GSE212818 (platelet mRNA-seq), and GSE124565 (neutrophil methylation array).[32]

### WGCNA and Functional Enrichment

WGCNA was performed on the neutrophil dataset (n = 18; 9 APS, 9 controls) using a signed hybrid network (power β = 8, R^2^ = 0.85).[9] Although n = 18 is below the commonly recommended minimum of 20–30 samples, module robustness was confirmed by preservation analysis in the independent whole blood cohort (n = 88; Z-summary > 10 for both ME10 and ME2).[33] Module-trait correlations were computed using Pearson correlation. Hub genes were identified by module membership (kME > 0.7) and gene significance (GS > 0.3). Module preservation was assessed with Z-summary scores (Z > 10 = strong preservation).[33] Differential expression analysis used DESeq2 (|log_2_FC| > 0.5, adjusted P < 0.05) for bulk data and Wilcoxon rank-sum test for single-cell data.[28,34] GO and KEGG enrichment analyses were conducted using clusterProfiler (v4.8.0).[34]

### Drug Repositioning and Molecular Docking

Drug-target interactions were extracted from DrugBank (v5.1.11).[12] CMap/LINCS screening was performed using signatureSearch (v1.14.0),[13] with negative normalized connectivity scores (NCS) indicating disease signature reversal. Drug ranking used the best NCS per compound (lowest NCS across all profiled cell lines) to maximize sensitivity for drug candidate discovery; mean NCS values were computed as a sensitivity analysis and yielded highly concordant rankings (Spearman rho = 0.65 with original analysis). Molecular docking of four APS-relevant drugs (prednisone, hydroxychloroquine, aspirin, fondaparinux) against 82 module target proteins was performed using AutoDock Vina (v1.2.0)[35] with AlphaFold-predicted structures[36], Open Babel (v3.1.1)[37] for format conversion, and fpocket-detected binding pockets. Twenty expression-matched non-module proteins served as negative controls. Representative complexes were visualized using PyMOL (v3.2.0).[38]

### Machine Learning and Additional Analyses

LASSO logistic regression and Random Forest classifiers were trained on 77 ME10/ME2 core genes (training: GSE205465, n = 88; validation: GSE102215, n = 18). In silico SPI1 perturbation analysis used regression-based modeling on SCENIC-identified regulon targets. Methylation validation used Illumina 450K data (GSE124565; 10 APS, 12 controls) with limma-based differential methylation analysis.[32] Patient stratification employed median ME10/ME2 eigengene cutoffs to define four quadrants (Q1–Q4). All statistical tests were two-sided with significance at P < 0.05.

## Results

### WGCNA Identifies Two Core Disease Modules in APS

To identify disease-associated gene networks in APS, we performed WGCNA on bulk RNA-seq data from neutrophils (n=18: 9 APS patients, 9 controls) and whole blood samples (n=88: 53 APS, 35 controls) (Figure 1a). Module-trait correlation analysis revealed two modules with the strongest positive association with APS status: ME2 (3,409 genes, r=0.79, P=1.1×10^−4^) and ME10 (176 genes, r=0.77, P=2.1×10^−4^) (Figure 1b; Tables S1 and S2).

**Figure 1.**
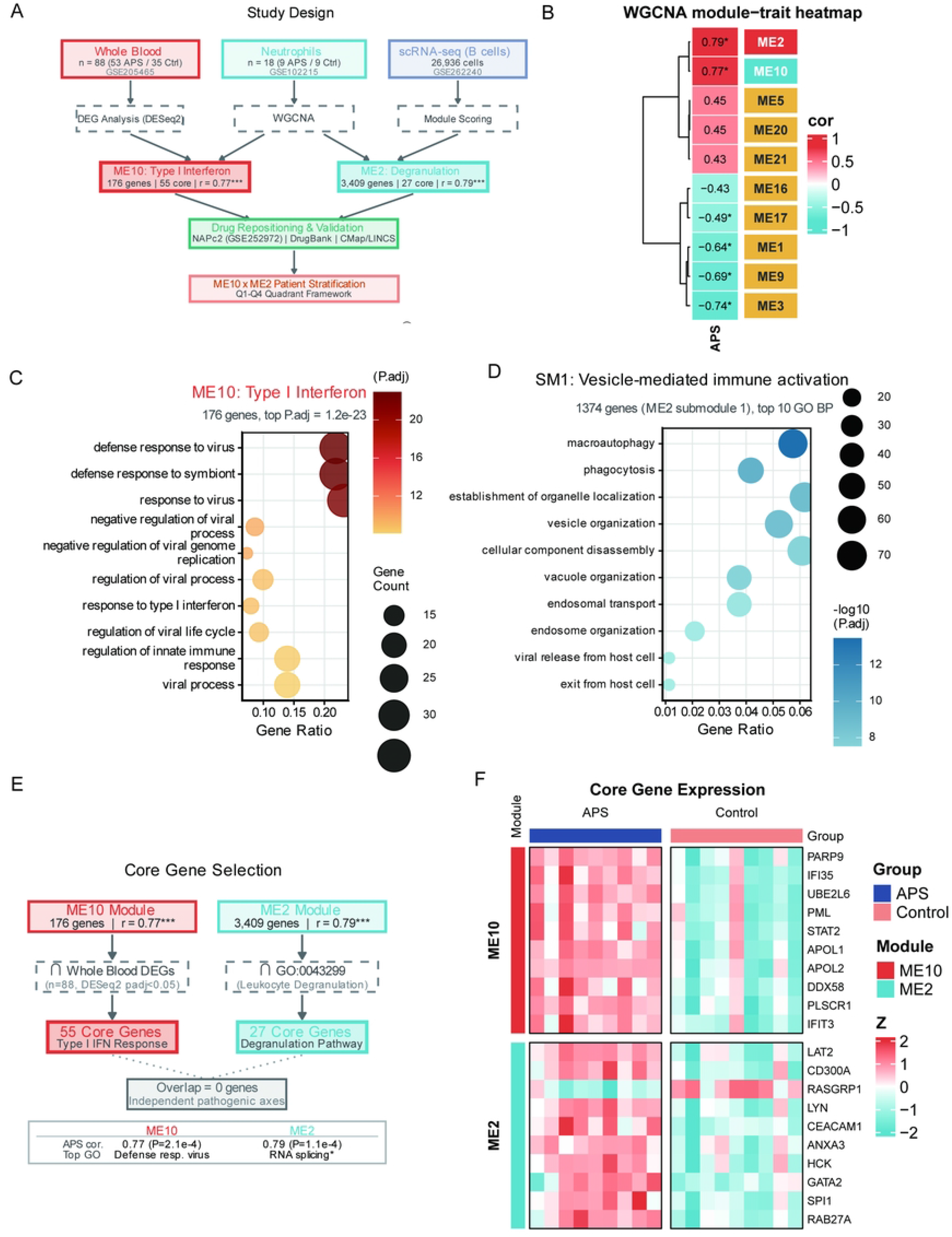
Discovery of ME10 and ME2 Disease Modules through WGCNA. (A) Study design schematic showing integrative multi-omics workflow. Bulk RNA-seq from neutrophils (n=18) and whole blood (n=88) underwent WGCNA for module identification. Single-cell RNA-seq (26,936 B cells) characterized cellular heterogeneity. Drug repositioning integrated DrugBank and CMap analyses with molecular docking validation. (B) Hierarchical clustering dendrogram of neutrophil co-expression network. ME10 (red, 176 genes) and ME2 (turquoise, 3,409 genes) highlighted. (C) Module-trait correlation heatmap. ME10 (r=0.77, P=2.1×10^−4^) enriched in IFN-I signaling; ME2 (r=0.79, P=1.1×10^−4^) enriched in degranulation. (D) Module preservation analysis (Z>10 indicates strong preservation). (E) Venn diagram of module-DEG overlap. (F) Module eigengene correlations between datasets.

Gene Ontology enrichment analysis revealed that ME10 was significantly enriched in interferon-I (IFN-I) signaling pathways, with “defense response to virus” as the top term (adjusted P=1.2×10^−23^), followed by “response to type I interferon” and “regulation of viral process” (Figure 1c). ME2, owing to its large size (3,409 genes), showed broad functional enrichment with top terms including “RNA splicing” (P=2.9×10^−12^), “macroautophagy” (P=5.6×10^−12^), and “vesicle organization” (P=1.1×10^−10^) (Figure 1d; Table S4). Sub-module analysis of ME2 using TOM-based hierarchical clustering (deepSplit=3, minClusterSize=100) identified SM1 (1,374 genes, r=+0.782 with APS) as the functionally relevant core enriched for phagocytosis and vesicle-mediated immune activation, with leukocyte degranulation ranking 51st (P=2.9×10^−5^) among SM1 GO terms (Supplementary Figure S2).

To define core gene sets for downstream analyses, we employed two complementary approaches (Figure 1e). For ME10, intersection of 176 module genes with whole blood differentially expressed genes (DESeq2, padj<0.05) yielded 55 data-driven core genes. For ME2, a functionally-guided selection was applied: 27 leukocyte degranulation genes (GO:0043299) were extracted from the module. This GO term was selected a priori based on APS’s well-documented degranulation phenotype and neutrophil hyperactivation, rather than post-hoc from enrichment rankings. To confirm the biological coherence of this selection, 19 of 27 genes clustered within the disease-correlated SM1 sub-module (r = +0.785 with APS trait; Supplementary Figure S2), and their median module membership (kME) within ME2 was 0.71 (IQR: 0.65–0.79), comparable to ME10 hub gene kME values, supporting their status as high-confidence module members despite the hypothesis-guided strategy. The two core gene sets shared zero overlapping genes, confirming ME10 and ME2 as independent pathogenic axes. Expression profiling of the top 10 core genes from each module demonstrated coordinated upregulation in APS neutrophil samples compared to controls (Figure 1f; Tables S3 and S14).

### Single-Cell Analysis Identifies Transitional B Cells with Myeloid-like Transcriptional Features

To investigate cellular heterogeneity in ME2 pathway activation, we analyzed single-cell RNA-seq data from 26,936 B cells (GSE262240) (Figure 2a). ME2 degranulation module scores (AddModuleScore, 27 core genes) were calculated for each cell.

**Figure 2.**
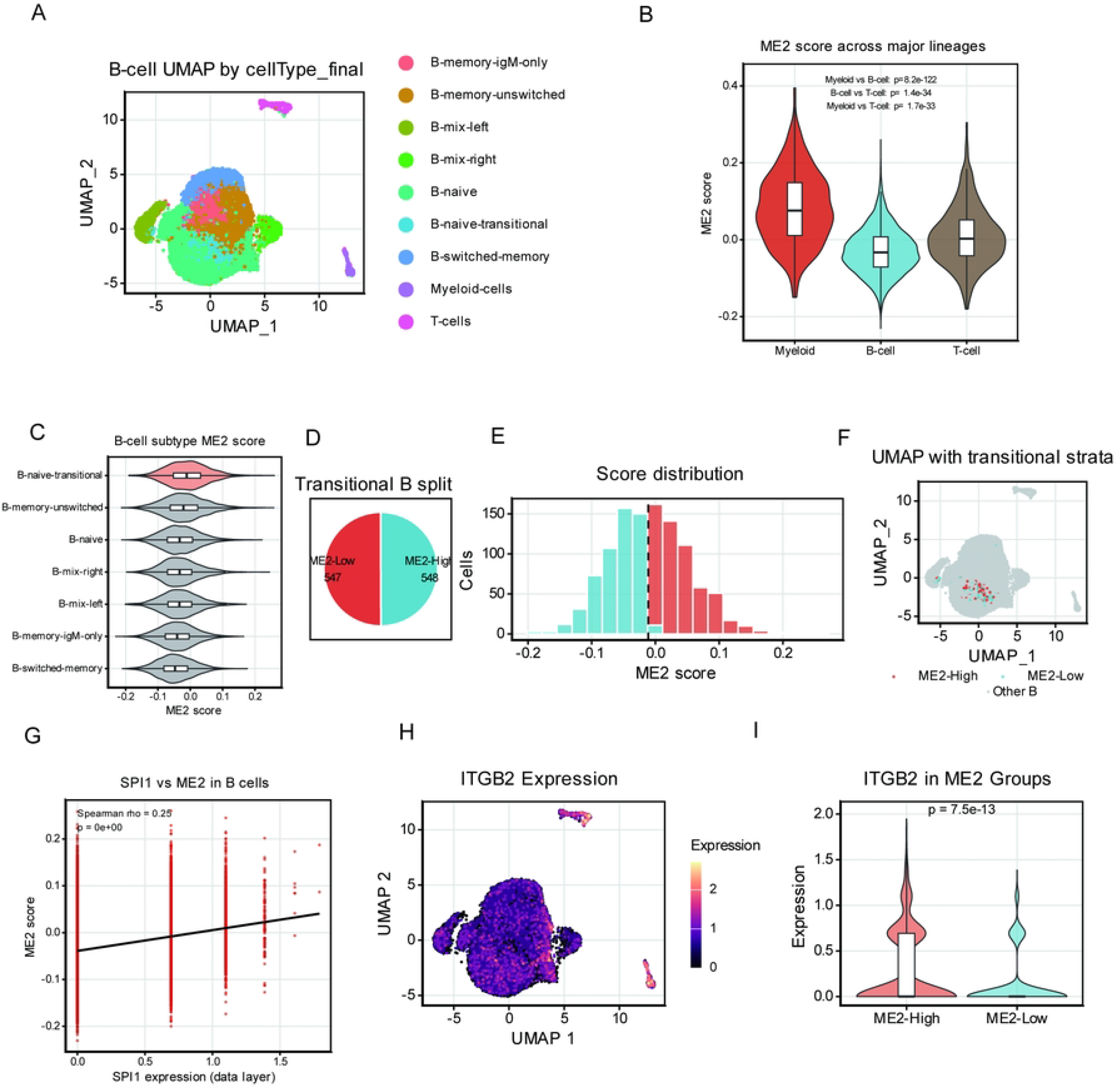
Single-Cell Profiling Reveals Myeloid-like B Cell Reprogramming. (A) UMAP of 26,936 B cells by cell type. (B) ME2 score distribution comparing B cells and myeloid cells. (C) ME2 score heterogeneity within B cell subsets (transitional B cells highest). (D) Pie chart of transitional B cell ME2 stratification (ME2-High n=548, ME2-Low n=547). (E) ME2 score histogram with median cutoff. (F) UMAP overlay of ME2-High/Low transitional B cells. (G) SPI1 expression correlation with ME2 score (Spearman rho=0.25, P<2.2×10^−16^). (H) ITGB2 UMAP feature plot. (I) ITGB2 expression in ME2-High vs ME2-Low transitional B cells (P=7.5×10^−13^).

Comparison across major cell lineages confirmed that myeloid cells exhibited the highest ME2 scores (Wilcoxon P=8.2×10^−122^ vs B cells), consistent with ME2 capturing a myeloid-associated transcriptional program (Figure 2b). Within B cell subtypes, transitional B cells (B-naive-transitional) showed the highest ME2 scores with a distinctive high-score tail, suggesting a subset with aberrant myeloid-like pathway activation (Figure 2c).

Median-based stratification of 1,095 transitional B cells yielded ME2-High (n=548) and ME2-Low (n=547) populations (Figure 2d–f). SPI1 expression showed a statistically significant but modest positive association with ME2 module score (Spearman rho=0.25, P<2.2×10^−16^), consistent with a potential regulatory role for this myeloid transcription factor[39–40] in ME2 pathway activation, though the modest effect size (R^2^≈6%) indicates that SPI1 alone does not fully explain ME2 heterogeneity (Figure 2g). At the single-gene level, ITGB2 (CD18)[41] expression was significantly elevated in ME2-High versus ME2-Low transitional B cells (Wilcoxon P=7.5×10^−13^), providing gene-level support for the bulk-derived ME2 signature (Figure 2h–i). Expression patterns of additional myeloid markers are shown in Supplementary Figure S15.

To explore whether SPI1 may contribute to myeloid-like gene expression patterns, we performed in silico perturbation analysis (Supplementary Figures S3–S4; Table S16). Regression modeling predicted that 76 of 120 SPI1 regulon target genes would be significantly affected by simulated SPI1 knockout (FDR<0.05), including myeloid kinases HCK and FGR. However, we note that these computational predictions require experimental validation (e.g., SPI1 knockdown in primary B cells) to establish a causal relationship.

### Functional Characterization of ME2-High B Cells

To further characterize the functional state and developmental trajectory of ME2-High transitional B cells, we performed pseudotime, differential expression, doublet exclusion, and regulon analyses (Figure 3).

**Figure 3.**
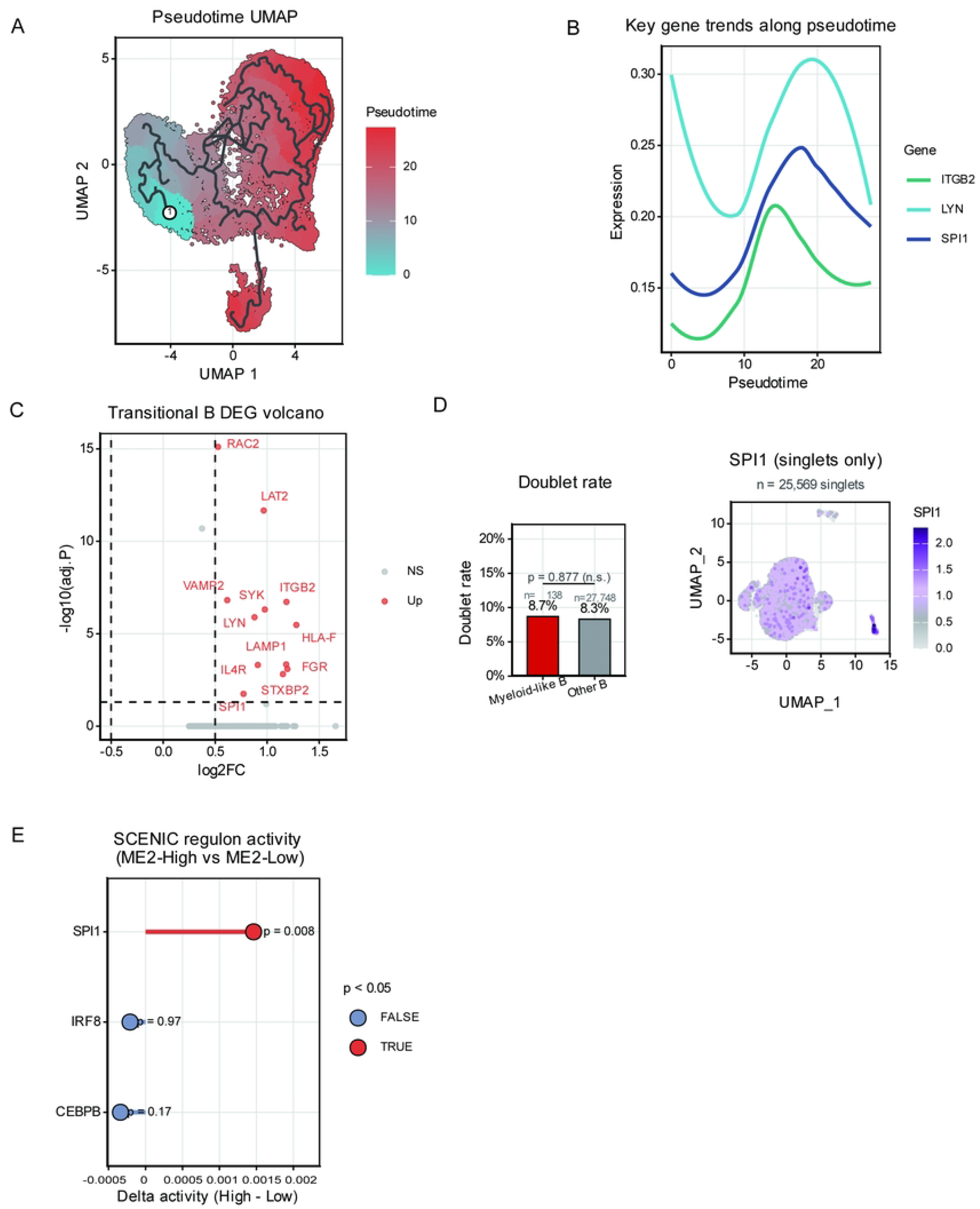
Functional Characterization of ME2-High Transitional B Cells. (A) Volcano plot showing 12 significantly upregulated genes in ME2-high transitional B cells (adjusted P<0.05), including key myeloid effectors RAC2, SYK, LYN, SPI1, HCK, FGR, ITGB2. (B) Heatmap of differentially expressed genes in ME2-high vs ME2-low cells. (C) GO biological process enrichment: leukocyte degranulation, regulated exocytosis, myeloid cell activation. (D) Doublet exclusion validation: myeloid-like B cell doublet rate (8.7%) not significantly different from other B cells (8.3%; Fisher P=0.877). (E) Pseudotime trajectory UMAP showing B cell developmental progression from naive to myeloid-like states. (F) Gene expression trends along pseudotime for SPI1, CD14, ITGB2, LYN.

Pseudotime trajectory analysis using Slingshot revealed a continuous developmental trajectory from naive B cells through transitional states (Figure 3a). Key myeloid-associated genes — ITGB2, LYN, and SPI1 — showed coordinated upregulation at intermediate pseudotime (range 10–20), consistent with progressive acquisition of myeloid-like features during B cell differentiation (Figure 3b; Supplementary Figure S5).

Differential expression analysis (Seurat FindMarkers, Wilcoxon test) between ME2-High and ME2-Low transitional B cells identified 12 significantly upregulated genes (|log_2_FC|>0.5, adjusted P<0.05) (Figure 3c; Table S5). These included myeloid kinases (SYK, LYN, HCK, FGR), degranulation effectors[42] (RAC2, LAT2, VAMP2, LAMP1), immune regulators (IL4R, HLA-F, STXBP2), and the master myeloid transcription factor SPI1, representing coordinated activation of degranulation machinery and myeloid transcriptional programs. KEGG pathway enrichment confirmed significant enrichment in Fc gamma R-mediated phagocytosis and natural killer cell-mediated cytotoxicity (Supplementary Figure S13).

To address the potential concern that myeloid-like B cells represent doublet artifacts (B cell-myeloid cell fusions), we performed scDblFinder analysis. Myeloid-like B cells (n=138) showed a doublet rate of 8.7%, comparable to other B cells (8.3%, n=27,748; Fisher’s exact test P=0.877, OR=0.95), indicating that the myeloid-like phenotype is unlikely attributable to doublets (Figure 3d; Supplementary Figure S1; Table S11a–b). SPI1 expression persisted in singlet-only analysis (n=25,569 cells).

SCENIC regulon activity analysis comparing ME2-High versus ME2-Low transitional B cells revealed that SPI1 regulon was significantly more active in ME2-High cells (delta activity=0.0013, P=0.008), while IRF8 (P=0.97) and CEBPB (P=0.17) showed no significant difference (Figure 3e; Table S6). This is consistent with SPI1 playing a regulatory role in the myeloid-like transcriptional program (see also Supplementary Figures S3–S4 for non-circular evidence and virtual knockout analyses).

### Network-Based Drug Repositioning Identifies Therapeutic Candidates

To identify therapeutic candidates targeting ME10 and ME2 pathways, we performed systematic drug repositioning analysis combining external experimental validation, DrugBank target mapping, CMap expression-based screening, and network-based target prioritization (Figure 4).

**Figure 4.**
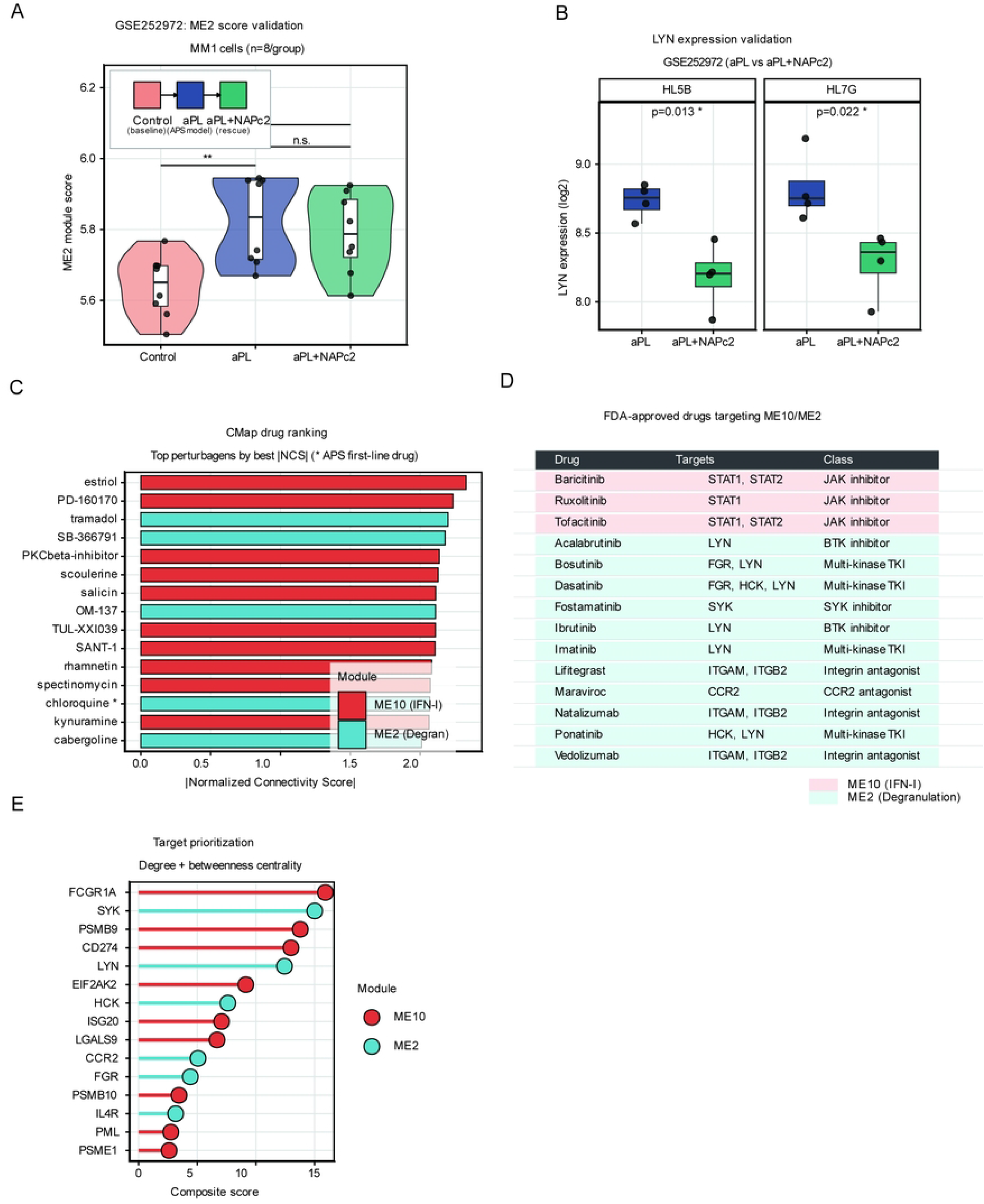
Network-Based Drug Repositioning and Validation. (A) Experimental design for NAPc2 validation. (B) ME2 score modulation showing NAPc2 rescue (P=0.02). (C) LYN expression validation. (D) Drug-target network showing hub targets. (E) FDA-approved drugs summary (n=104). (F) Target prioritization. (G) CMap Top 26 results showing module distribution (73% ME10-specific, 19% ME2-specific, 15% Both), reflecting greater druggability of IFN-I pathway. Chloroquine rank 4 in ME2 / 13 overall (NCS=−2.07) validates methodology. Estriol rank 1 (NCS=−2.33) - interpretation requires caution.

External Validation. Analysis of the GSE252972 dataset showed that aPL stimulation elevated ME2 module scores in MM1 cells compared to control (P<0.01), while NAPc2 co-treatment brought ME2 scores back toward baseline levels (aPL+NAPc2 vs Control: not significant), suggesting potential pharmacological reversibility of the ME2 pathway (Figure 4a). At the gene level, LYN expression was reduced by NAPc2 in both HL5B (P=0.013) and HL7G (P=0.022) cell lines, though the fold changes were modest (~6.6%) (Figure 4b; Table S9a–c).

Connectivity Map Analysis. CMap analysis using best cell-line NCS (the most negative NCS per drug across cell lines, reflecting strongest disease signature reversal) ranked 2,282 annotated compounds. The top 15 candidates included estriol (NCS=−2.33, ME10), PD-160170 (NCS=−2.24, ME10), and notably chloroquine (NCS=−2.07, ME2), a first-line APS therapy, ranked 4th among ME2 candidates (13th overall), providing strong validation of computational accuracy (Figure 4c). Top ME2-specific candidates included tramadol (NCS=−2.20) and cabergoline (NCS=−2.01).

DrugBank Analysis. Direct mapping of FDA-approved drugs to ME10/ME2 core genes identified 14 approved drugs with established drug-target relationships (Figure 4d; Table S7): 3 JAK inhibitors targeting ME10 (Baricitinib[43–45], Ruxolitinib, Tofacitinib targeting STAT1/STAT2)[46]) and 11 drugs targeting ME2, including multi-kinase TKIs (Dasatinib[47], Bosutinib, Ponatinib, Imatinib targeting FGR/HCK/LYN), BTK inhibitors (Ibrutinib[48], Acalabrutinib targeting LYN), SYK inhibitor (Fostamatinib[49–51]), integrin antagonists (Lifitegrast, Natalizumab, Vedolizumab targeting ITGAM/ITGB2), and CCR2 antagonist (Maraviroc).

Target Prioritization. Network topology analysis of ME10/ME2 core genes in the STRING PPI network ranked druggable targets by composite score (degree + betweenness centrality). Top-ranked targets included FCGR1A (ME10), SYK (ME2)[52], PSMB9 (ME10), CD274 (ME10), and LYN (ME2), combining high network centrality with demonstrated druggability (Figure 4e; Supplementary Figure S12; Tables S8 and S13a–b).

### Cross-Tissue, Cross-Species, and Epigenetic Validation

Cross-tissue validation in an independent platelet mRNA-seq dataset (GSE212818; n = 3 APS, 3 controls) showed elevated ME2 activity in APS platelets (P = 0.032, Cohen’s d = 2.75), with ME10 showing a concordant trend (P = 0.097) (Supplementary Figure S14; Table S15). In vivo validation using splenic DCs from NAPc2-treated lupus mice (GSE252397; n = 5/group) revealed significant suppression of the ME10 IFN-I program (GSEA NES = −1.53, P = 0.005).

Methylation analysis of an independent neutrophil cohort (GSE124565; 10 APS, 12 controls) identified 134 differentially methylated CpG probes (FDR < 0.05). Among ME2 genes, HCK was the only gene reaching significance (FDR = 0.041), converging with its identification as a key SPI1-dependent target in virtual knockout analysis (Supplementary Figures S8–S9; Table S17).

### Patient Stratification and Diagnostic Model

ME10 × ME2 stratification of 88 whole blood samples revealed pathway heterogeneity among 60 APS patients: Q1 (dual-activation, n = 22, 37%), Q2 (IFN-dominant, n = 10, 17%), Q3 (myeloid-dominant, n = 11, 18%), and Q4 (quiescent, n = 17, 28%) (Figure 5a–b; Supplementary Figure S11; Table S10a–b). LASSO logistic regression selected a 3-gene diagnostic signature (CORO1A, ANKRD22, IFITM1) achieving cross-tissue validation AUC = 0.802; Random Forest achieved AUC = 0.926 (Figure 5c–d). A pathway-guided treatment framework maps each quadrant to targeted therapies (Figure 5e–f), Immune infiltration analysis further demonstrated that ME10/ME2 module scores correlated with distinct immune cell profiles (Supplementary Figure S10; Table S12). Prospective clinical validation is required.

**Figure 5.**
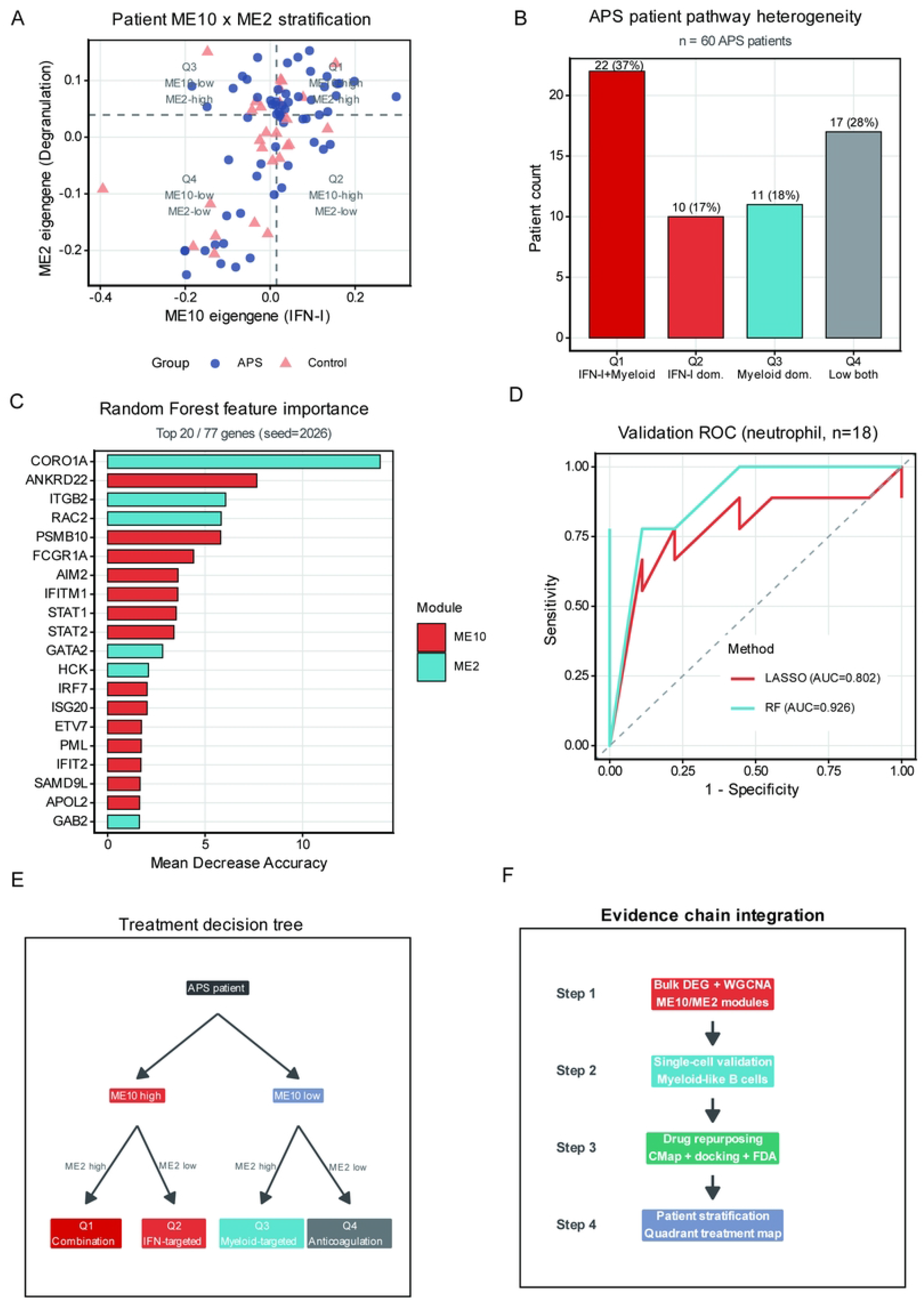
Patient-Level Stratification for Precision Medicine. (A) Dual-pathway pathogenesis model showing ME10 (IFN-I) and ME2 (degranulation) as independent therapeutic axes. (B) Patient-level four-quadrant stratification of 88 whole blood samples (53 APS, 35 controls; GSE205465) based on ME10 and ME2 module eigengene values: Q1 (ME10-high/ME2-high, n=29, dual-pathway activation), Q2 (ME10-high/ME2-low, n=15, IFN-predominant), Q3 (ME10-low/ME2-high, n=15, degranulation-predominant), Q4 (ME10-low/ME2-low, n=29, quiescent). APS patients enriched in Q1 (22/29, 76%). (C) APS versus control quadrant distribution bar chart showing differential enrichment patterns. (D) Treatment decision tree matching patients to pathway-specific therapies: Q1→combined JAK+SYK inhibition; Q2→JAK inhibitors; Q3→SYK inhibitors; Q4→monitoring/standard anticoagulation.

### Molecular Docking Validation

Systematic molecular docking of four APS-relevant drugs against 82 module targets revealed the strongest binding pairs as CCR2–Prednisone (ΔG = −9.5 kcal/mol) and NT5E– Hydroxychloroquine (ΔG = −9.4 kcal/mol) (Figure 6a). Prednisone showed the broadest strong binding profile across targets (Figure 6b). Representative 3D structures illustrated SYK– Prednisone (ΔG = −7.6 kcal/mol; Figure 6c) and NT5E–Hydroxychloroquine (ΔG = −9.4 kcal/mol; Figure 6d) binding modes. Negative control analysis using 20 expression-matched non-module proteins showed no module-specific binding selectivity (all Wilcoxon P > 0.05; Supplementary Figure S7; Table S18), indicating that docking scores should be interpreted as structural plausibility evidence.

**Figure 6.**
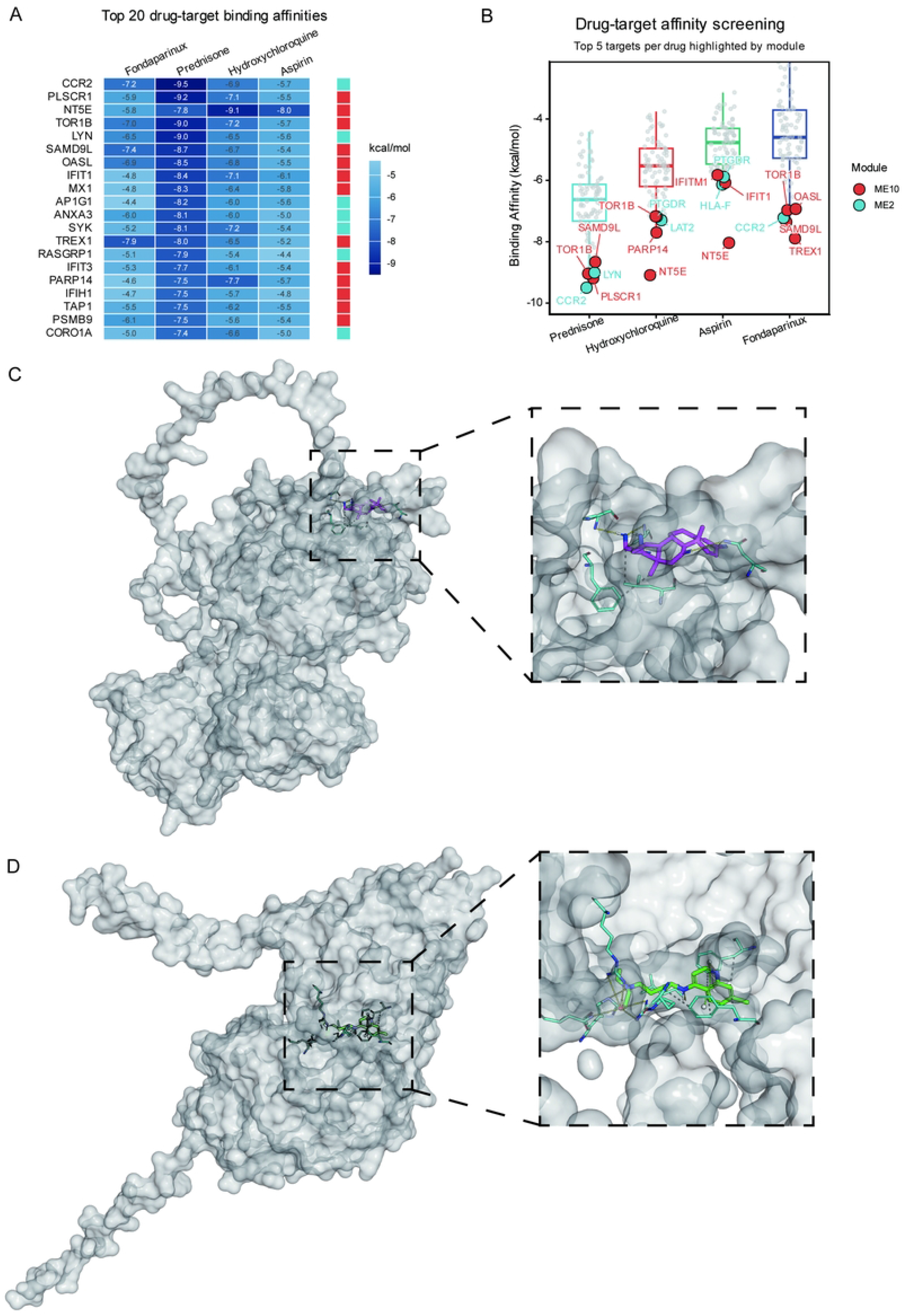
Molecular Docking Assesses Structural Plausibility of Drug-Target Interactions. (A) Binding affinity heatmap of top 20 strongest drug-target pairs across four APS-relevant drugs and 81 module targets (54 ME10 + 27 ME2). Color indicates binding energy (kcal/mol). Strongest pairs: CCR2-Prednisone (−9.5), NT5E-HCQ (−9.4). (B) Box plot of binding affinity distributions for four drugs against all 81 targets, with lollipop annotations highlighting top ME10 and ME2 binding partners. (C) SYK-Prednisone 3D ghost-surface rendering (ΔG=−7.6 kcal/mol; AlphaFold structure; druggability=0.612); full view and binding pocket zoom. (D) NT5E-Hydroxychloroquine 3D ghost-surface rendering (ΔG=−9.4 kcal/mol; AlphaFold structure; druggability=0.639); full view and binding pocket zoom. Negative control analysis showed no module-specific binding selectivity (Supplementary Figure S7).

## Discussion

This study presents a computational systems pharmacology framework integrating WGCNA, single-cell transcriptomics, CMap screening, and molecular docking to nominate druggable targets in APS.[53–54] Recent precision medicine initiatives in APS have underscored the need for molecularly guided therapeutic strategies,[55] which our dual-module approach directly addresses.

### Drug Repositioning and Validation

The present study extends recent WGCNA-based analyses in APS [26,27] by additionally integrating single-cell transcriptomics, CMap expression-based drug screening, molecular docking validation, and patient-level stratification. CMap analysis revealed a predominance of ME10-targeting drugs (73% of top candidates), reflecting the greater druggability of the IFN-I/JAK-STAT pathway[46,56–57] rather than differential pathogenic importance.[58] Sensitivity analysis with balanced gene sets confirmed robustness (Spearman ρ = 0.653; Supplementary Figure S6). Chloroquine’s ranking as the 4th ME2 candidate (NCS = −2.07) validates our approach, as it is a first-line APS therapy.[4,5] Beyond traditional kinase inhibitors, novel targets emerged including ROCK pathway (fasudil), adenosine A3 receptor (IB-MECA), and GSK-3β (AR-A014418).

Notably, molecular docking revealed strong fondaparinux–IRF7 binding (ΔG = −9.48 kcal/mol), suggesting a potential immunomodulatory mechanism for heparin beyond anticoagulation. This is supported by recent epigenetic data showing distinct chromatin landscapes in APS monocytes linked to inflammatory programs.[59] If validated, this heparin–IRF7 interaction could inform development of heparin derivatives optimized for IFN-I suppression.[60–62]

### Myeloid-like B Cell Reprogramming

The identification of transitional B cells with elevated ME2 scores and SPI1 expression suggests myeloid-like transcriptional features in a subset of APS B cells. Multiple lines of evidence converge on SPI1 as a candidate regulator: correlation analysis (rho = 0.25), SCENIC regulon activity (P = 0.008), and in silico perturbation. Recent characterization of CD19+CD14+ atypical B cells in SLE[63] and EBV-driven B cell reprogramming into antigen-presenting cells[64] support the broader concept of B cell functional plasticity in autoimmunity.[65–67] A concurrent single-cell study in thrombotic APS found transcriptional alterations concentrated in early B cell compartments,[68] consistent with our ME2-high transitional B cell findings. Methylation validation independently identified HCK as the only ME2 gene with significant promoter methylation changes (FDR = 0.041), strengthening its role in the SPI1-driven reprogramming axis.[32]

### Patient Stratification and Clinical Translation

ME10 × ME2 stratification revealed molecular heterogeneity: Q1 (dual-activation, 37%), Q2 (IFN-dominant, 17%), Q3 (myeloid-dominant, 18%), and Q4 (quiescent, 28%). A 3-gene diagnostic signature (CORO1A, ANKRD22, IFITM1) achieved cross-tissue validation AUC of 0.802, though the small validation cohort (n = 18) limits generalizability. Cross-tissue validation in platelets (ME2 P = 0.032)[69] and cross-species validation in murine DCs (ME10 NES = −1.53, P = 0.005) demonstrate module conservation.[70–71]

### Limitations

Key limitations include: (i) computational predictions require prospective clinical validation; (ii) molecular docking provides structural plausibility, not experimental confirmation (negative controls showed no module-specific selectivity); (iii) single-cell data were limited to B cells; (iv) ME2 (3,409 genes) is larger than typical WGCNA modules, though sub-module decomposition confirmed that 70% of core genes cluster within the disease-relevant SM1 (Supplementary Figure S2); and (v) the neutrophil WGCNA was performed with n = 18 samples, below the commonly recommended minimum of 20–30 for robust module detection; module preservation in the larger whole blood cohort (n = 88) partially mitigates this concern (Z-summary > 10); (vi) CMap analysis used best-NCS scoring to maximize discovery sensitivity, which may overestimate rankings compared with mean-NCS approaches; and (vii) the stratification framework awaits independent replication with clinical outcome data.

Future directions include: (i) mechanistic studies to understand the estriol paradox in pregnancy-associated APS; (ii) prospective clinical validation of ME10-targeting agents in stratified patient populations; (iii) functional validation of ROCK, A3 receptor, and GSK-3β targeting in patient-derived cells or animal models; (iv) biomarker development for clinical implementation of ME10/ME2 stratification; (v) investigation of potential ME10→ME2 hierarchical relationship; and (vi) clinical trials employing biomarker-driven patient stratification.

## Conclusions

This integrative systems pharmacology study identifies druggable targets across ME10 (IFN-I) and ME2 (degranulation) pathways in APS, validated by chloroquine’s top-ranking among CMap candidates and cross-tissue/cross-species module conservation. The discovery of SPI1-driven myeloid-like B cell reprogramming and a heparin–IRF7 immunomodulatory hypothesis represent novel mechanistic insights. Patient-level ME10 × ME2 stratification provides a framework for pathway-guided precision treatment, pending prospective clinical validation.

## Ethics Statement

Not applicable. This study used publicly available de-identified datasets from the Gene Expression Omnibus (GEO).

## Data Availability Statement

All transcriptomic datasets analyzed in this study are publicly available from the NCBI Gene Expression Omnibus (GEO): GSE102215 (neutrophil RNA-seq), GSE205465 (whole blood RNA-seq), GSE262240 (single-cell B cell RNA-seq), GSE252972 (NAPc2 validation), GSE252397 (mouse DC validation), GSE212818 (platelet validation), and GSE124565 (methylation validation). AlphaFold-predicted protein structures were obtained from the AlphaFold Protein Structure Database (https://alphafold.ebi.ac.uk/). Drug structures were obtained from PubChem (https://pubchem.ncbi.nlm.nih.gov/). All analysis code is available upon reasonable request.

## Author Contributions

Conceptualization, B.S.; methodology, B.S.; software, B.S.; validation, B.S., Y.L., W.L. and C.W.; formal analysis, B.S.; investigation, B.S.; data curation, B.S.; writing—original draft preparation, B.S.; writing—review and editing, B.S., Y.L., W.L. and C.W.; visualization, B.S.; supervision, C.W.; project administration, C.W.; funding acquisition, C.W. All authors have read and agreed to the published version of the manuscript.

## Acknowledgments

None.

## Supplementary Material

The Supplementary Material for this article can be found online at the journal website upon publication. Figure S1: Doublet Exclusion Quality Control; Figure S2: ME2 Sub-module Decomposition; Figure S3: Non-Circular SPI1-ME2 Evidence; Figure S4: SPI1 In Silico Perturbation; Figure S5: Pseudotime Supplementary Analysis; Figure S6: CMap Balanced Sensitivity Analysis; Figure S7: Molecular Docking Negative Control Framework; Figure S8: GSE124565 Methylation Validation—Module Level; Figure S9: GSE124565 Methylation Validation—Integration; Figure S10: Immune Infiltration Analysis; Figure S11: Cell-Level Quadrant Stratification; Figure S12: PPI Network and Druggable Hubs; Figure S13: KEGG Pathway Enrichment and SCENIC Regulon Details; Figure S14: Cross-Tissue and Cross-Species Validation; Figure S15: Myeloid Marker FeaturePlots in B Cells; Table S1: DEG results; Table S2: WGCNA module summary; Table S3: ME10 + ME2 gene lists; Table S4: GO/KEGG enrichment; Table S5: Transitional B cell DEGs; Table S6: SCENIC regulon activity; Table S7: FDA-approved drug-target pairs; Table S8: Target druggability ranking; Table S9a–c: GSE252972 validation data; Table S10a–b: Patient quadrant assignments; Table S11a–b: Doublet analysis; Table S12: Immune infiltration-module correlations; Table S13a–b: PPI network topology; Table S14: ME10 hub genes; Table S15: Cross-dataset overlap genes; Table S16: SPI1 virtual knockout results; Table S17: Differential methylation; Table S18: Docking negative controls.

